# Mutations in the microexon splicing regulator *srrm4* have minor phenotypic effects on zebrafish neural development

**DOI:** 10.1101/2024.11.29.626094

**Authors:** Tripti Gupta, Gennady Margolin, Harold A. Burgess

## Abstract

Achieving a diversity of neuronal cell types and circuits during brain development requires alternative splicing of developmentally regulated mRNA transcripts. Microexons are a type of alternatively spliced exon that are 3–27 nucleotides in length and are predominantly expressed in neuronal tissues. A key regulator of microexon splicing is the RNA-binding protein Serine/arginine repetitive matrix 4 (*Srrm4*). *Srrm4* is a highly conserved, vertebrate splicing factor that is part of an ancient family of splicing proteins. To better understand the function of *Srrm4* during brain development, we examined neural expression of zebrafish *srrm4* from days 1–5 of development using fluorescence *in situ* hybridization. We found that *srrm4* has a dynamically changing expression pattern, with expression in diverse cell types and stages during development. We then used CRISPR-based mutagenesis to generate zebrafish *srrm4* mutants. Unlike previously described morphant phenotypes, *srrm4* mutants did not show overt morphological defects. Moreover, sequencing of wild-type and mutant transcriptomes revealed only minor changes in splicing. *srrm4* thus appears to have a limited role in zebrafish neural development.

## Introduction

The diversity of cell types and intricate circuitry of vertebrate brains is the result of complex developmental processes that occur during embryonic and early post-embryonic stages. These processes include specification of neuronal precursors, differentiation into distinct cell types, axonal migration, and synaptogenesis, with regulation of these processes occurring at both a spatial and temporal level. Forward genetic screens as well as targeted mutagenesis studies identified many of the genes required for neural development. More recently, large scale transcriptomic studies and advances in bioinformatic analyses have revealed a significant cell type-specific diversity in transcript variants, highlighting the importance of alternative splicing during neural development (Weyn-Vanhentenryck et al. 2018; Furlanis et al. 2019; Booeshaghi et al. 2021; Leung et al. 2021; Patowary et al. 2024).

Alternative splicing allows the generation of multiple protein isoforms from a single gene, resulting in increased molecular diversity. This increased diversity facilitates the development of tissues such as the brain, in which there are many different cell types that communicate through complex patterns of connectivity. Sequencing studies have shown that alternative splicing occurs at a high frequency in neural tissues and that it is highly conserved (reviewed in Raj and Blencowe 2015). Microexons are a particular class of alternatively spliced exon that are 3–27 nucleotides in length (reviewed in Ustianenko et al. 2017 and Gonatopoulos-Pournatzis and Blencowe 2020). Microexon inclusion most often preserves the reading frame of the protein, and microexons are predicted to encode amino acids in protein-protein interaction domains (Irimia et al. 2014). One study found that neural-enriched microexons are more highly conserved than any other class of microexon, indicating their importance in neural development and function (Irimia et al. 2014).

A key regulator of microexon splicing in the nervous system is the serine/arginine repetitive matrix 4 gene (*Srrm4*; also named *nSR100*), which has been shown to regulate inclusion of over 50% of profiled microexons (Irimia et al. 2014). This vertebrate-specific splicing regulator belongs to the SR family of alternating Arg/Ser domain-containing proteins that function in spliceosome assembly (Calarco et al. 2009). *In vitro* studies have shown that neurite outgrowth is impaired in Neuro2a cells when *Srrm4* levels are knocked down (Calarco et al. 2009; Ohnishi et al. 2017). Mice homozygous for a C-terminal deletion of SRRM4 display increased anxiety and deficits in hearing and balance due to defective hair cell differentiation (Nakano et al. 2012; Shirakawa et al. 2024). A second allele of mouse *Srrm4*, with a frame-shifting exon deletion that results in loss of the full length protein, has more severe phenotypes that include disrupted neurite outgrowth and cortical layering, as well as autistic-like behavioral deficits (Quesnel-Vallières et al. 2015; Quesnel-Vallières et al. 2016). Accordingly, it was found that individuals with autism spectrum disorder had lower levels of *SRRM4* mRNA and a corresponding misregulation of neural microexons, demonstrating the potential importance of this gene in neurodevelopmental disorders (Irimia et al. 2014).

While most studies of *Srrm4* function have been limited to two loss-of-function mouse mutants and *in vitro* models, two studies in zebrafish utilizing morpholinos demonstrated effects of *srrm4* knockdown on neuronal differentiation, axonal arborization, and hair cell development (Calarco et al. 2009; Nakano et al. 2012). However, the spatiotemporal pattern of *Srrm4* expression during neural development has not been described in detail, and there are concerns about the reliability of even well-controlled morpholino studies (reviewed in Stainier et al. 2015, Lawson 2016, and Arana and Sánchez 2024). We therefore investigated the functions of *srrm4* during zebrafish brain development by first performing a detailed characterization of its expression from 1 to 5 days post fertilization (dpf). We then established a mutant model to determine effects on brain morphology and function, as well as to enable identification of additional *srrm4* microexon targets. Zebrafish are ideal for studies of neural development because the skin is translucent during embryonic and early larval stages, enabling imaging of the developing brain. In addition, established techniques for whole brain morphometric analysis of brain structure and composition allow unbiased screening for phenotypes affecting all regions of the brain (Randlett et al. 2015; Gupta et al. 2018), in contrast to previous studies of *Srrm4*, which have focused on candidate brain regions.

We found that *srrm4* is broadly expressed throughout the brain during development but also has distinct areas of high expression. Through co-localization with *notch1a* and *elavl3*, we determined that *srrm4* is expressed in proliferating and newly differentiated neurons, depending on the region and developmental stage. Notably, there was a high level of *srrm4* expression in the granule cells of the cerebellum and the torus longitudinalis, suggesting a role for *srrm4* in granule cell development. We then used a whole brain morphometric approach to look for changes in neuronal composition and region-specific differences in brain volume. We found that knockdown of *srrm4* in G0 crispant larvae resulted in a decrease in optic tectum neuropil. Unexpectedly, stable mutant alleles did not show this phenotype. We sequenced mRNA from *srrm4* mutants and sibling wild-type animals and detected only minor differences in alternative splicing of microexons. The absence of strong phenotypes in *srrm4* stable mutants suggested the presence of genetic compensation; however, RNA-Seq analysis uncovered only a very small number of differentially expressed genes, none of which revealed an obvious mechanism for genetic compensation. It is also possible that redundancy with the *srrm4* paralog *srrm3* may mask mutant phenotypes. Thus, our analysis of *srrm4* has identified brain regions and cell types in which microexon splicing is potentially important; however, it will be necessary to overcome potential redundancy and/or compensation to better understand the functions of *srrm4* during neural development.

## Materials and Methods

### Animal husbandry

Experiments were conducted in accordance with institutional guidance and approved by the National Institute of Child Health and Human Development Animal Care and Use Committee. All lines were maintained in a Tüpfel Longfin (TL) background. Larvae were reared at 28° C with a 14 hour light/10 hour dark cycle. For all imaging experiments except 24 hpf HCR staining and FM1-43 hair cell staining, larvae were raised in 300 µM N-Phenylthiourea in E3 (Sigma catalog # 222909) beginning at 8-24 hpf to inhibit melanogenesis.

### *in situ* hybridization chain reaction (HCR)

Larvae were fixed for 2 hours at room temperature or overnight at 4° C in 4% paraformaldehyde (EMS catalog # 15710) and 4% sucrose (MP Biomedicals catalog # 802536) in 1X phosphate buffered saline (PBS; catalog # AM9625), washed 4 x 20 minutes in 1X PBS, dehydrated in 100% methanol, and stored at -20° C for at least 24 hours. For 2–5 dpf larvae, skin was sometimes dissected away from the brain after fixation in PFA. HCR was performed according to the protocol in (Choi et al. 2018) with the following modifications: proteinase K was not used for permeabilization and the larvae were not post-fixed. B1, B2, and B3 amplifiers were purchased from Molecular Instruments, as were all other HCR reagents. *elavl3* probe was purchased from Molecular Instruments. All other probe pools were designed using HCR 3.0 Probe Maker software (Kuehn et al. 2022) and purchased from Integrated DNA Technologies (IDT oPools™). Probe pool sequences are in Supplementary Table 4.

### Immunofluorescence

Larvae were fixed overnight at 4° C in 4% paraformaldehyde (EMS catalog # 15710) in 1X phosphate buffered saline (PBS; catalog # AM9625), washed 4 x 20 minutes in 1X PBS, dehydrated in 100% methanol, and stored at -20° C. Larvae were rehydrated to 1X PBST (1X PBS with 0.01% Tween) and then blocked for two hours in blocking solution (5% NGS, 1% BSA, and 1% DMSO in 1X PBST) at room temperature. Samples were then incubated overnight at 4° C in primary antibody solution: mouse anti-acetylated Tubulin (1:500; Sigma catalog # T7451) and mouse anti-Islet-1 40.2D6 (1:500; Developmental Studies Hybridoma Bank) diluted in blocking solution. After 2 rinses and 3 twenty minute washes in 1 X PBST, samples were incubated overnight at 4° C in secondary antibody solution: Alexa Fluor 488 goat anti-mouse IgG1 (1:500; Invitrogen A-21121) and Alexa Fluor 546 goat anti-mouse IgG2b (1:500; Invitrogen A-21143) diluted in blocking solution. Following staining, larvae were washed at least 3 times for 20 minutes each and then mounted in Vectashield (Fisher catalog # NC9265087).

### Image acquisition and analysis

Confocal images were acquired using a Nikon A1R inverted scanning confocal microscope using 20X and 40X water immersion objectives and 488 nm, 561 nm, and 647 nm laser lines. Fixed samples were mounted in 50% glycerol/50% 5 x SSC (Quality Biological catalog # 351-003-101). Live imaged samples were anesthetized in 0.2% tricaine methanesulfonate (Sigma catalog # E10521) in E3 embryo medium and imaged in 2.5% low melt agarose (Lonza catalog # 50080) in E3 as described (Bhandiwad et al. 2024).

Image processing was done using Fiji software (Schindelin et al. 2012). Images were registered and averaged using Advanced Normalization Tools (ANTs) (Avants et al. 2009 Jul 29; Avants et al. 2011) using previously described parameters (Marquart et al. 2017) run on the NIH Biowulf computing cluster.

Morphometric analyses of G0 crispant and stable mutant larvae at 6 dpf were conducted using ANTs for brain registration and CobraZ software for statistical analysis, as described in (Gupta et al. 2018). Brains were registered using one or more transgenes or with lysotracker dye as described in (Bhandiwad et al. 2024). Transgenic lines used were: *vglut2a:GFP* (derived from *TgBAC(slc17a6b:lox-DsRed-lox-GFP)nns14* by Cre injection) (Satou et al. 2013); *gad1b:RFP* (*TgBAC(gad1b:lox-RFP-lox-GFP)nns26 (gad1b*:*RFP*)) (Satou et al. 2013), and *tuba:mCar (Tg(-2.6Cau.tuba1:mCar.zf1)y516* (*tuba:mCar*)) (Gupta et al. 2018). For details on specific experiments, see Table 1. gRNA cutting in crispants was confirmed with CRISPR-STAT genotyping (Carrington et al. 2015). Primer sequences are available in Supplementary Table 6.

**Table 1:** Descriptions of experimental conditions for morphometry experiments described in text and Figure 7. Sequence descriptions of mutants and crispants are in Supplementary Table 5. Mut: mutant; het: heterozygote; wt: wild-type.

|  |  |
| --- | --- |
| **G0 crispant experiments** |  |
| 1 | Crispant A and Cas9/tracr control in *vglut2a:GFP; gad1b:RFP; tuba:mCard* background |
| 2 | Crispant A and Cas9/tracr control in *Gal4<sup>y304</sup>* background (Gal4 strongly expressed in optic tectum) |
| 3 | Crispant B and Cas9/tracr control in *vglut2a:GFP* background |
| 4 | Crispant C and Cas9/tracr control in *tuba:mCard* background |
| 5 | Crispant D and Cas9/tracr control in *tuba:mCard* background |
| **Stable mutant experiments** | **Genotypes Tested and Conditions** |
| 1 | *srrm4<sup>y713</sup>* mut and wt from het incross in *vglut2a:GFP; tuba:mCard; gad1b:RFP* background |
| 2 | *srrm4<sup>y712</sup>* mut and wt from het incross stained with lysotracker |
| 3 | *srrm4<sup>y712</sup>* mut and wt from het incross injected with Cas9/tracr stained with lysotracker |
| 4 | *srrm4<sup>y712</sup>* mut and wt "cousins" stained with lysotracker |
| 5 | *srrm4<sup>y712</sup>* mut and wt from het incross in *vglut2a:GFP; tuba:mCard* background |
| 6 | *srrm4<sup>y712</sup>* mut female x het male and *srrm4<sup>y712</sup>* wt female x wt male in *tuba:mCard* background |

### Statistical analyses

Repeated measures ANOVA was performed using JASP software (JASP Team (2024). Meta-analysis was conducted with PyMeta software using a random effects model and mean difference effect measure (Deng, H., PythonMeta).

### FM1-43 staining

3 dpf larvae were incubated in 3 µM FM™ 1-43 Dye (Thermofisher catalog # T3163) for 2-4 minutes, then anesthetized and mounted in low melt agarose for imaging, as described above.

### gRNA microinjections

4.6 nl of gRNA/sgRNA+Cas9 protein injection mix was injected into each 1 cell TL or transgenic embryo using a Nanoject II Microinjector (Drummond Scientific). Embryos were then reared in E3 embryo medium at 28° C with a 14 hour light/10 hour dark cycle. *srrm4* and *atoh1c* Alt-R™ CRISPR-Cas9 crRNA (gRNA targets are listed in Supplementary Table 6) were ordered from Integrated DNA Technologies (IDT). Alt-R® CRISPR-Cas9 tracrRNA and Alt-R™ S.p. Cas9 Nuclease were also purchased from IDT (Catalog #s 11-01-03-01 and 1081058). Some experiments were done using a *srrm4* target 4 sgRNA (Supplementary Table 6) from Synthego (CRISPRevolution sgRNA EZ Kit) with Cas9 Nuclease from IDT.

### Generation of stable mutations

gRNAs were designed using ChopChop (Labun et al. 2019). No off-target effects were predicted to be in *srrm3* or any other known splicing factor. *srrm4^y712^* was generated by injecting 600 pg total of four target gRNAs + 767 pg of Cas9 protein into TL embryos. Cutting with three of the four gRNAs (targets 1, 2, and 3) was confirmed by CRISPR-STAT genotyping (Carrington et al. 2015); gRNA target and primer sequences in Supplementary Table 6). *srrm4^712^* was generated by injecting 600 pg of *srrm4* target 4 gRNA + 767 pg of Cas9 protein into TL embryos. F1 embryos from G0 adults were screened for cutting and raised to adulthood. Mutations in F1 adults were identified by CRISPR-STAT genotyping and outcrossed to TL fish to generate stable mutant lines. Mutations in F2 adults were confirmed by Sanger Sequencing (Supplementary Table 5).

### G0 crispant knockdowns

Descriptions of *srrm4* crispant experiments are provided in Table 1. The same gRNAs used in crispant experiments were used to make stable mutant lines (see above for details). For crispant experiments 1 and 2, 600 pg total of four target gRNAs + 767 pg of Cas9 protein was injected into each embryo. For crispant experiment 3, 900 pg total of two target gRNAs + 4700 pg of Cas9 protein was injected per embryo. For crispant experiment 4, 450 pg of target 4 gRNA + 4700 pg of Cas9 protein was injected. For crispant experiment 5, 1000 pg of target 5 gRNA + 4700 pg of Cas9 protein was injected. For the *atoh1c* C0 crispant, gRNAs were designed using ChopChop (Labun et al. 2019); Supplementary Table 6). 900 pg total of two target gRNAs + 4700 pg of Cas9 protein was injected per embryo.

### Protein alignments

Protein alignments were performed using Jalview and Clustal W and Muscle (Edgar 2004; Waterhouse et al. 2009). Percent identities were calculated using JalView. Zebrafish eMIC domain was defined based on InterProScan of Pfam database and sequence match to Pfam15230 domain.

### RNA-Seq

*srrm^y712^* heterozygous adults were incrossed to obtain larvae for sequencing. At 3 dpf, larvae were tail-clipped for CRISPR-STAT genotyping and the heads were stored in RNAlater™ (Invitrogen catalog # AM7024) at 4° C. After identification of homozygous mutant and homozygous wild-type larvae, 8–10 larvae per genotype were pooled and total RNA was isolated using the Qiagen RNeasy mini kit (Qiagen catalog # 74104) or Zymo Direct-zol RNA Microprep Kit (Zymo Research catalog # R2060). 2 x 100-bp paired-end sequencing was performed on 1 µg of total RNA by the NICHD Molecular Genomics Core facility. RNA-Seq was performed on three biological replicate samples. RNA-Seq data for all six samples is available in the NCBI Sequence Read Archive database using accession # PRJNA1189474.

### DESeq2 analysis

We used DESeq2 v1.34.0 for standard differential gene expression analysis of RNA-Seq data from 3 dpf *srrm4^y712^* homozgygous mutant and homozygous wild-type samples (Love et al. 2014). Three biological replicates were used, and each mutant sample was paired with the sibling wild-type sample from the same heterozygous incross (design *∼batch+genotype*.). Supplementary Table 2 contains *log2FoldChange* and *lfcSE* values estimated through the “ashr” shrinkage procedure (which does not affect p-values) (Stephens 2017).

### Vast-Tools analysis

Vast-Tools analysis was performed on RNA-Seq data from the 3 dpf *srrm4^y712^* homozgygous mutant and homozygous wild-type samples described above. We downloaded VAST-tools (Tapial et al. 2017) v2.2.2 and zebrafish database file *vastdb.dre.01.12.18.tar.gz* (https://github.com/vastgroup/vast-tools). We ran *vast-tools* sub-commands with default parameters for paired-end sequencing, and ran vast-tools diff to generate a file of differential analysis between the two conditions. This file was filtered to select MV[dPSI] scores over 10% as well as *DreEX* events (alternatively spliced exons for *Danio rerio*). We then used the VASTDB database (https://vastdb.crg.eu/wiki/Main_Page) to determine the sizes of the exons, which were added as the final column. This yielded the 59 entries listed in Supplementary Table 1.

## Results

### Zebrafish *srrm4*

Zebrafish contains a single *srrm4* gene on chromosome 5 with two predicted transcript variants. The longer variant (NCBI Reference Sequence NM_001423807.1), which we will refer to as transcript 1, encodes a predicted 574 amino acid protein (NP_001410736.1) that is 44% identical to human NP_919262.2 and 47% identical to mouse NP_001390326.1 (Figure 1A and B). Near the C-terminus of the protein, zebrafish isoform 1 contains the conserved eMIC domain that is reported to be required and sufficient for microexon splicing when expressed in HEK293 cells (Torres-Méndez et al. 2019). Within the eMIC domain, the zebrafish and mouse sequences are 64% identical, indicating a high degree of conservation. The shorter variant (transcript 2, XM_021476416.1) encodes a predicted 254 amino acid protein. This variant contains a 16 nt microexon close to the 3’ end of the transcript that is similar to that found in mouse *Srrm4* (Quesnel-Vallières et al. 2016; ENSMUST00000222119.2). Inclusion of this microexon causes a frame-shift that is predicted to result in a premature stop codon, leading to a truncated protein lacking the functional eMIC domain (Quesnel-Vallières et al. 2016).

**Figure 1:**
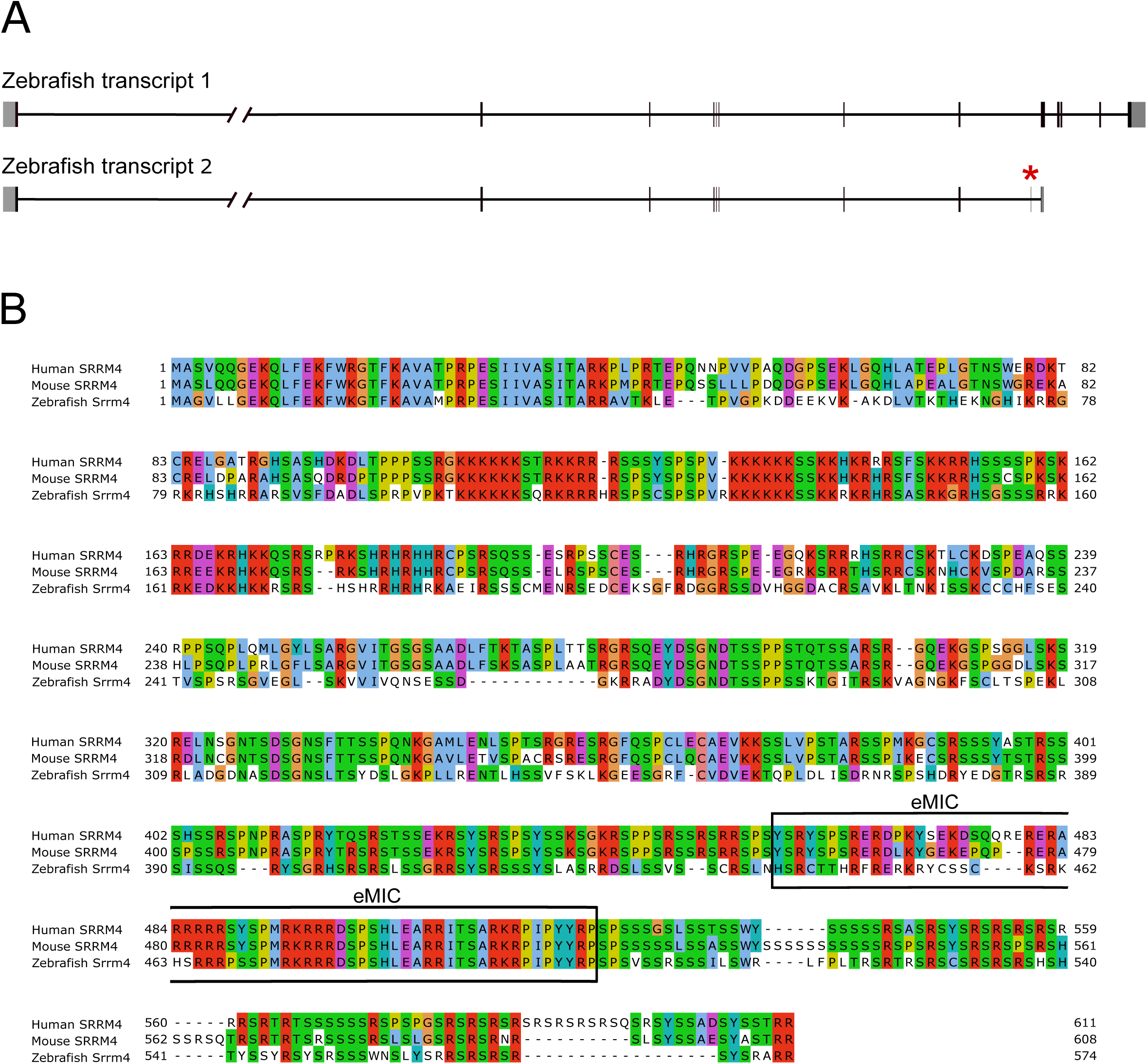
Zebrafish *srrm4* has two annotated transcript variants and encodes a protein with a conserved eMIC domain. (A) *srrm4* transcript variants. Transcript variant 1 (NM_001423807.1) encodes the longer isoform of the Srrm4 protein, which contains the eMIC domain. Transcript variant 2 (XM_021476416.1) has a 16 nt microexon near the 3’ end that creates a frame-shift, resulting in a truncated protein lacking the eMIC domain. Red asterisk indicates microexon. Grey shading indicates untranslated regions. (B) Sequence alignment of longest isoforms of human, mouse, and zebrafish SRRM4/Srrm4 proteins. Black box outlines the conserved eMIC domain.

### *srrm4* is dynamically expressed during neural development

The expression pattern of *srrm4* from 24–48 hpf was previously reported by (Calarco et al. 2009). We performed high resolution fluorescence *in situ* hybridizations for *srrm4* expression from 24 hours post fertilization (hpf) to 5 dpf, using confocal microscopy and co-localization with cell type-specific markers to more precisely define cellular expression. At 24 hpf, we observed *srrm4* expression in the telencephalon and olfactory epithelia of the forebrain, the diencephalon, the tegmentum, and throughout the developing hindbrain (Figures 2A and 2B), as well as more ventral structures (data not shown). At 48 hpf, *srrm4* is expressed at high levels in regions that include the optic tectum cellular layer, developing cerebellum, and hindbrain (Figures 2C and 2D). Within the hindbrain, *srrm4* exhibits a striped pattern, typical of rhombomeric expression. To determine if *srrm4* expression is restricted to a specific cell state, we first tested co- localization of *srrm4* and *notch1a*, a marker for proliferating cells (Mueller and Wullimann 2016). At 48 hpf, *srrm4* and *notch1a* strongly co-localize in the hindbrain, indicating expression of *srrm4* in proliferating cells of the rhombomeres (Figures 2E–E’’, (Belmonte-Mateos et al. 2023). Interestingly, rhombomere boundary cells also show partial co-localization of *srrm4* with *elavl3*, a marker for newly differentiated cells (Mueller and Wullimann 2016) (Figures 2F–F’’). By 3 dpf, *srrm4* is expressed in regions throughout the neuraxis (Figures 3A and 3B). In some regions, including the optic tectum cellular layer, the medial habenula and a portion of the torus semicircularis, there was strong co-expression of *srrm4* and *elavl3* (Figures 3A’’ and 3B’’). However, in other regions, such as within the mesopontine tegmentum, there was high expression of *srrm4* and low expression of *elavl3* (Figure 3B’’). At 4 dpf (not shown) and 5 dpf, *srrm4* remains broadly expressed, but notable differences in cell-type expression are present. In the cerebellum, *srrm4* and *elavl3* show significant co-localization in the middle of the corpus cerebelli, but only partially co-localize in the lobus caudalis (Figures 3C–C’’). The varying co-localization of *srrm4* with *notch1a* and *elavl3* in different cell types shows that *srrm4* expression does not delineate a specific cell state during neural development.

**Figure 2:**
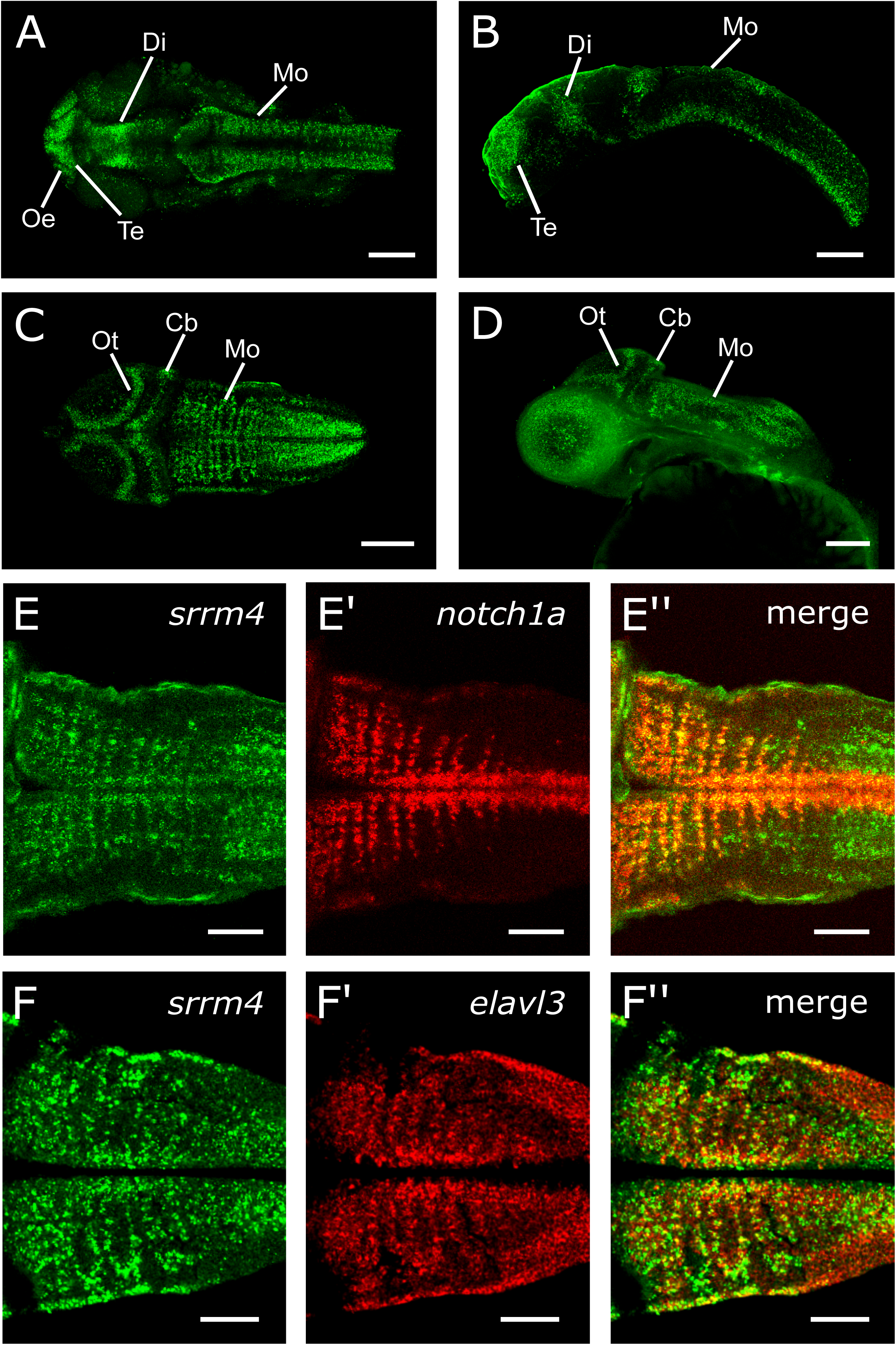
*srrm4* is expressed in proliferating *notch1a* and differentiated *elavl3* neurons during early brain development. (A-D) Partial (20 µm depth) maximum intensity projections of *srrm4* mRNA expression at 24 hpf and 48 hpf. Dorsal view in (A) and sagittal view in (B) at 24 hpf. Dorsal view in (C) and sagittal view in (D) at 48 hpf. Anterior is to the left. Scale bars = 100 µm. (E-E’’) Single plane image of Mo of *srrm4* co- labeled with *notch1a* mRNA at 48 hpf. (E) *srrm4* expression. (E’) *notch1a* expression. (E’’) *srrm4* and *notch1a* merged images. (F-F’’) Single plane image of Mo of *srrm4* co-labeled with *elavl3* mRNA at 48 hpf. (F) *srrm4* expression. (F’) *elavl3* expression. (F’’) *srrm4* and *elavl3* merged images. Scale bars = 50 µm. Cb: cerebellum; Di: diencephalon; Mo: medulla oblongata; Oe: olfactory epithelium; Ot: optic tectum; Te: telencephalon.

**Figure 3:**
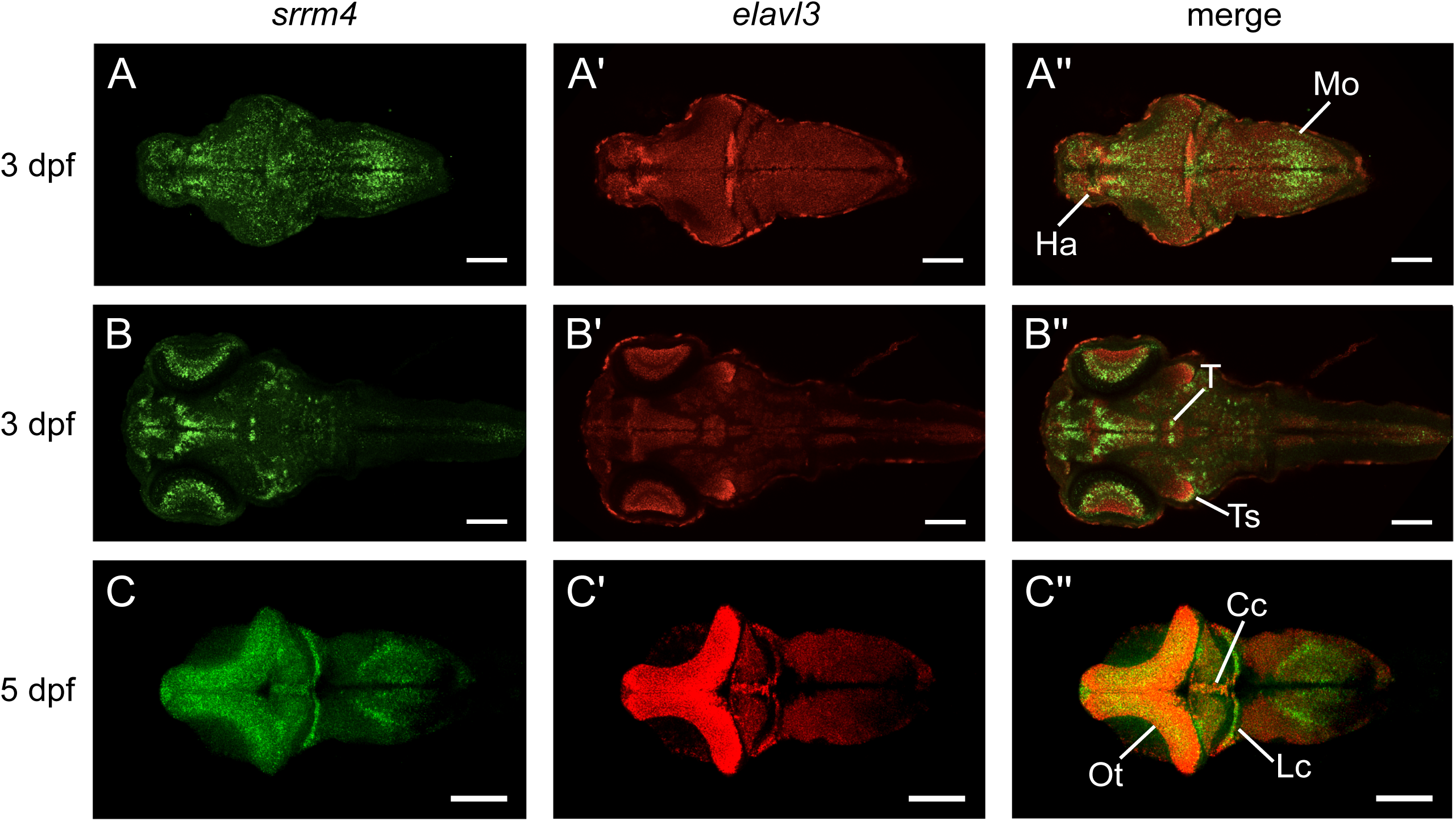
*srrm4* is dynamically expressed at 3 dpf and 5 dpf. (A–B’’) Single plane images of *srrm4* and *elavl3* mRNA expression at 3 dpf in two focal planes (3 brains averaged). Dorsal views of *srrm4* (A and B), *elavl3* (A’ and B’), and *srrm4* and *elavl3* merged (A’’ and B’’). (C–C’’) *srrm4* and *elavl3* mRNA expression at 5 dpf (5 brains averaged). Dorsal views of *srrm4* (C), *elavl3* (C’), and *srrm4* and *elavl3* merged (C’’). Anterior is to the left. Scale bars = 100 µm. Cc: Corpus cerebelli; Ha: habenula; Lc: lobus caudalis; Mo: medulla oblongata; Ot: optic tectum; T: tegmentum; Ts: torus semicircularis.

In examining the expression of *srrm4* in the hindbrains of 3–5 dpf larvae, we observed that *srrm4* was expressed in longitudinal columns that extended along the rostro-caudal axis of the brain (Figure 4). This striped expression pattern was similar to that described by (Kinkhabwala et al. 2011) for glutamatergic and glycinergic neurons. To investigate if *srrm4* labels cells of a specific neurotransmitter type in the developing hindbrain, we performed *in situ* hybridization for *srrm4* and *glyt2* to label glycinergic neurons and *srrm4* in a *vglut2a-GFP* transgenic background to label glutamatergic neurons at 4 dpf. We found very limited co-localization of *srrm4* and *vglut2a-GFP* that was restricted to the dorsal aspects of columns in the caudal medulla oblongata (Figure 4A–B’’). In contrast, we detected more significant but still only partial co-localization of *srrm4* and *glyt2* (Figure 4C–D’’). Similarly, *srrm4* partially co-localized with *gad1b* in the hindbrain (data not shown). Therefore, despite its longitudinal expression in the developing hindbrain, *srrm4* does not demarcate neurons of a particular neurotransmitter type.

**Figure 4:**
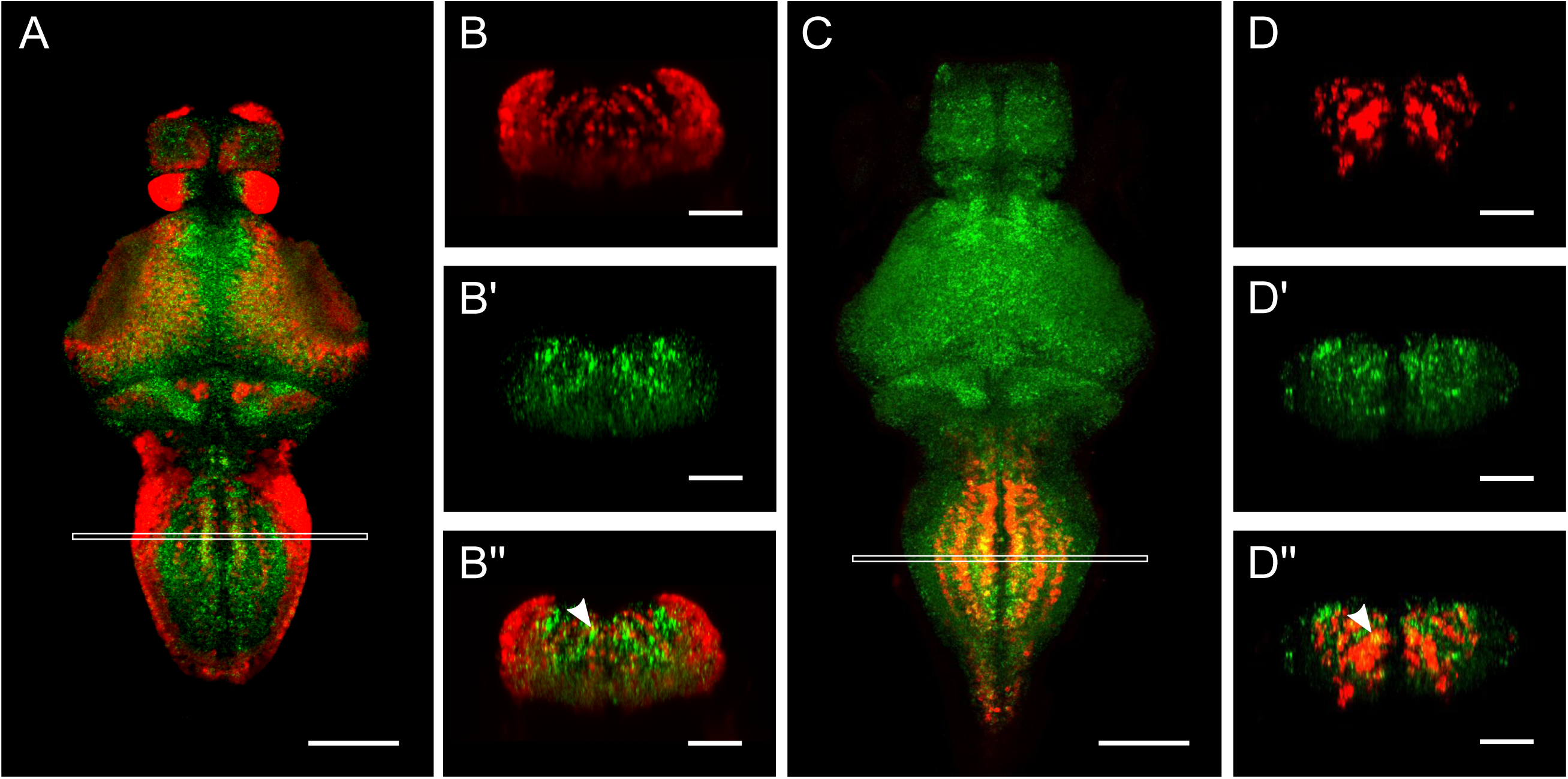
*srrm4* partially co-localizes with glutamatergic and glycinergic neuron markers in the medulla. (A–B’’) *srrm4* mRNA and *vglut2a:GFP* transgene expression at 4 dpf. (A) Dorsal view of single plane image showing *srrm4* and *vglut2a:GFP* expression (5 brains averaged). White box indicates area of maximum intensity projection shown in B-B’’ from a single representative brain. Coronal views of *vglut2a:GFP* (B), *srrm4* (B’), and *vglut2a:GFP* and *srrm4* merged (B’’). (C–D’’) *srrm4* mRNA and *glyt2* expression at 4 dpf. (C) Dorsal view of single plane image showing *srrm4* and *glyt2* expression (5 brains averaged). White box indicates area of maximum intensity projection shown in D-D’’ from a single representative brain. Coronal views of glyt2 (D), *srrm4* (D’), and glyt2 and *srrm4* merged (D’’). Scale bars in A and C = 100 µm. Scale bars in B–B’’ and D–D’’ = 50 µm. Arrowheads point to examples of co-localization.

### *srrm4* is expressed in granule cell precursors

Expression of *srrm4* in the developing cerebellum is especially striking and of particular interest, as high levels of *srrm4* expression have been reported in the mouse cerebellum, as well (Nakano et al. 2012; Nakano et al. 2020). We noted that expression of *srrm4* in the cerebellum is highest in the corpus cerebelli, eminentia granularis and lobus caudalis, all of which contain large numbers of glutamatergic granule cells. In addition, we found that *srrm4* is expressed at high levels in the torus longitudinalis, a cerebellar-like structure unique to ray-finned fishes (chondrosteans, holosteans and teleosts) that is also predominantly composed of granule cells. To test whether *srrm4* is expressed in granule cells, we performed *in situ* hybridization chain reaction (HCR) for *srrm4* and the granule cell marker *vglut1* (Bae et al. 2009). We found that at 6 dpf, *srrm4* colocalizes with *vglut1* in the lobus caudalis (Fig. 5A–A’’), eminentia granularis (data not shown), and corpus cerebelli (Fig. 5B–B’’) of the cerebellum. In addition, *srrm4* colocalizes with *vglut1* in the torus longitudinalis (Fig. 5C–C’’). These data confirm that *srrm4* is expressed in multiple populations of *vglut1* expressing granule cells.

**Figure 5:**
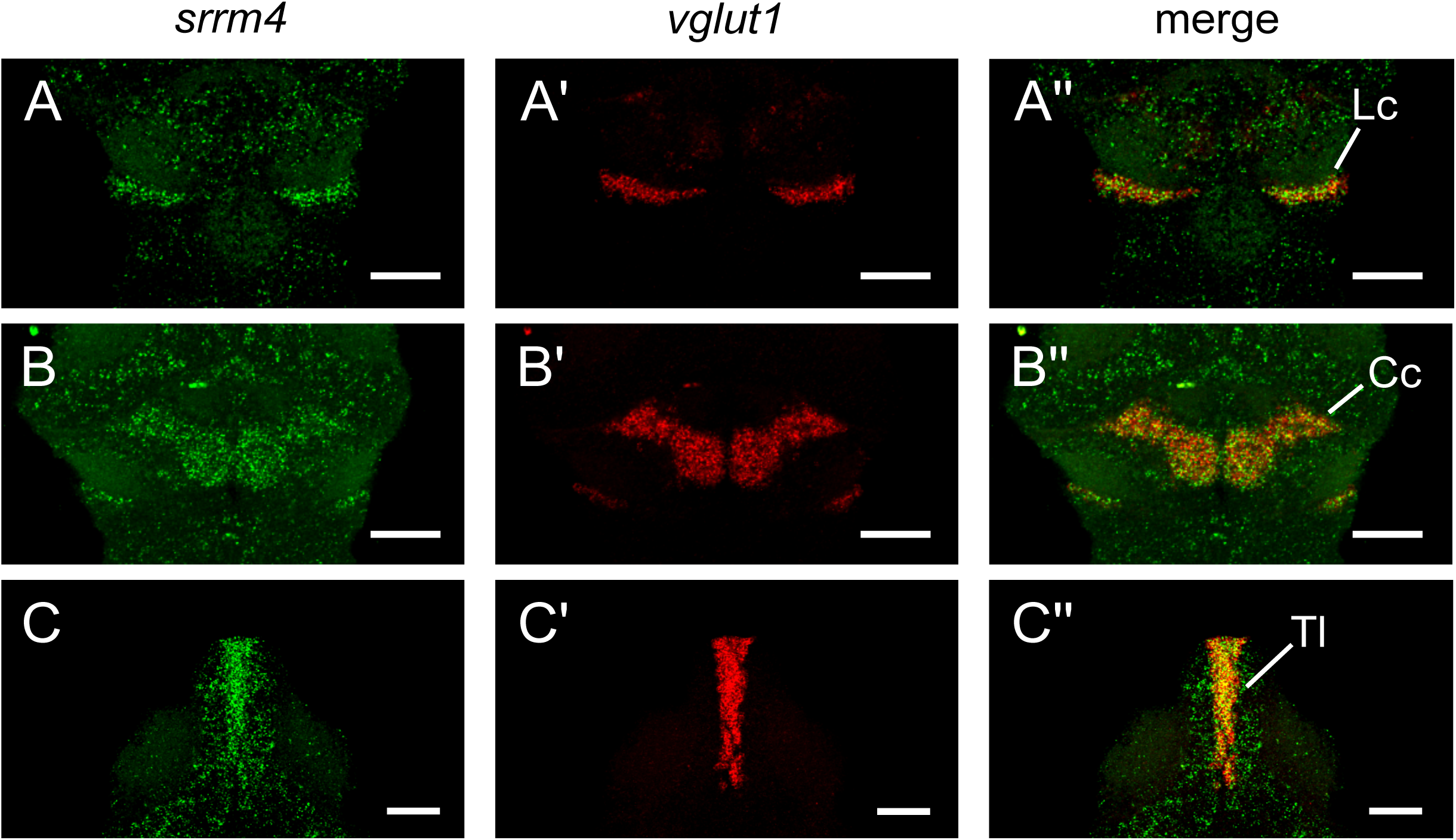
*srrm4* co-localizes with the granule cell marker *vglut1*. (A–C’’) *srrm4* and *vglut1* mRNA expression at 6 dpf. (A–A’’) Dorsal view of the lobus caudalis showing *srrm4* (A), *vglut1* (A’), and *srrm4* and *vglut1* merged (A’’). (B–B’’) Dorsal view of the corpus cerebelli showing *srrm4* (B), *vglut1* (B’), and *srrm4* and vglut1 merged (B’’). (C–C’’) Dorsal view of the torus longitudinalis showing *srrm4* (C), *vglut1* (C’), and *srrm4* and *vglut1* merged (C’’). Anterior is to the top. Scale bars = 50 µm. Cc: Corpus cerebelli; Lc: lobus caudalis; Tl: torus longitudinalis.

In mammals, the granule cells of the cerebellum develop within a region of the hindbrain called the upper rhombic lip (URL) from cells that express the transcription factor Atoh1 (Ben-Arie et al. 1997). Similarly, granule cells in the zebrafish cerebellum develop from cells in the URL that migrate rostrally to the corpus cerebelli and valvula cerebelli, laterally to the eminentia granularis, and to the lobus caudalis (Volkmann et al. 2008; Kani et al. 2010). While granule cell development in mammals is regulated by a single *Atoh1* gene, zebrafish have three *atoh1* orthologs termed *atoh1a*, *atoh1b*, and *atoh1c*. *atoh1a* and *atoh1b* are initially expressed in the URL starting at 24 hpf, with expression becoming largely localized to the valvula cerebelli by 5 dpf (Kaslin et al. 2009; Kani et al. 2010; Kidwell et al. 2018). *atoh1c* expression is first detected in the URL at 2 dpf and by 3 dpf is found in corpus cerebelli, lobus caudalis, and eminentia granularis cells (Kidwell et al. 2018). We hypothesized that *srrm4* expression in the granule cell precursors may be regulated by one or more *atoh1* genes. Since the majority of granule cells are specified by *atoh1c*, we tested co-localization of the *srrm4* and *atoh1c* transcripts by *in situ* hybridization from 2 dpf to 4 dpf. At 4 dpf, *srrm4* and *atoh1c* expression domains show significant overlap within the URL (Fig. 6A–A’’). By 3 dpf, *srrm4* transcripts show a broader expression domain than *atoh1c* that extends rostrally from the URL (Fig. 6B–B’’). We interpret this as expression of *srrm4* in the developing granule cells as they migrate to the corpus cerebelli, valvula cerebelli, and lobus caudalis. By 4 dpf, *atoh1c* and *srrm4* are largely expressed in distinct domains with *atoh1c* expression in the URL and *srrm4* in the corpus cerebelli (not shown) and lobus caudalis (Figure 6C–C’’). In an *atoh1c* G0 crispant knockdown, expression of *srrm4* was not detected in the lobus caudalis and was reduced in the corpus cerebelli (Supplementary Figure 1). These data suggest that *srrm4* expression is downstream of *atoh1c* in the URL.

**Figure 6:**
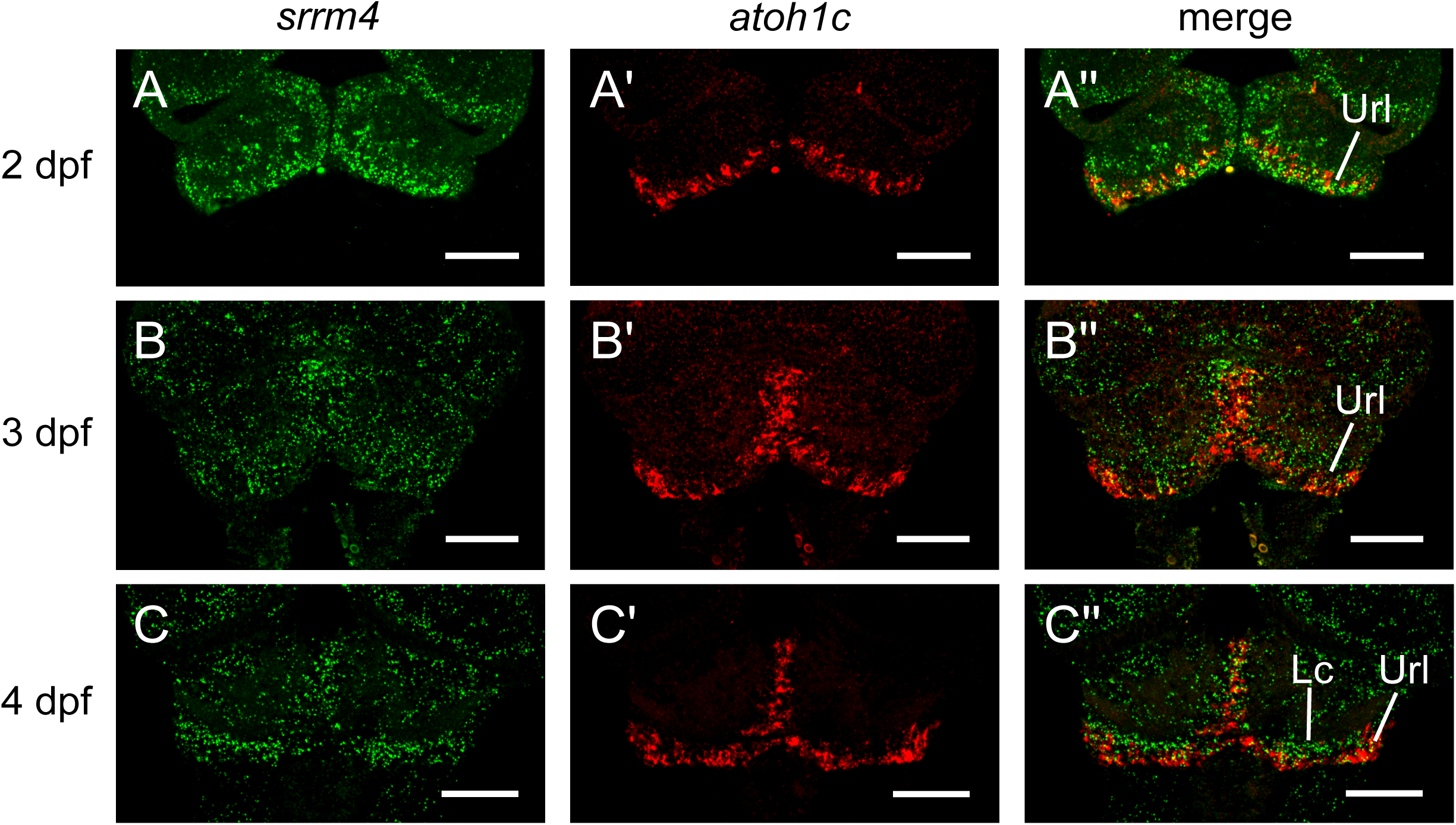
*srrm4* co-localizes with *atoh1c* in the Url. (A–C’’) *srrm4* and *atoh1c* mRNA expression at 2–4 dpf. (A–A’’) Partial maximum intensity projection of upper rhombic lip at 2 dpf showing *srrm4* (A), *atoh1c* (A’), and *srrm4* and *atoh1c* merged (A’’). (B–B’’) Partial maximum intensity projection of upper rhombic lip at 3 dpf showing *srrm4* (B), *atoh1c* (B’), and *srrm4* and *atoh1c* merged (B’’). (C–C’’) Partial maximum intensity projection of upper rhombic lip and lobus caudalis at 4 dpf showing *srrm4* (C), *atoh1c* (C’), and *srrm4* and *atoh1c* merged (C’’). Dorsal views. Anterior is to the top. Scale bars = 50 µm. Lc: lobus caudalis; Url: upper rhombic lip.

### *srrm4* mutant lines do not recapitulate morphant phenotypes

In order to study the *in vivo* function of *srrm4,* we generated two mutant lines using CRISPR/Cas9-mediated mutagenesis. For *srrm4^y712^*, we targeted four independent sites to increase our chances of generating a non-functional allele. Three of the four guide RNAs (gRNAs) were effective, resulting in a 14 nucleotide (nt) deletion in exon 1, a 3 nt deletion and single nt change in exon 6, and a 5 nt deletion in exon 10. Frameshifts caused by these deletions result in a predicted truncated protein that is 48 amino acids long (Supplementary Figure 2A). Because the conserved eMIC domain was shown to be necessary for *srrm4* function *in vitro*, we generated a second allele designed specifically to disrupt this domain. We generated *srrm4^y713^* by using a single gRNA that targets the eMIC domain and screening for mutations that would produce in-frame deletions to avoid potentially triggering nonsense-mediated decay and subsequent genetic compensation. *srrm4^y713^* has a 9 nt deletion and four single nucleotide changes in the eMIC domain, resulting in a protein lacking three amino acids and with three amino acid changes (Supplementary Figure 2B). Homozygous mutants of both alleles were viable to adulthood and were fertile.

We next assessed mutants for loss-of-function phenotypes, starting with those previously reported for *srrm4* morphants. At 30 and 36 hpf, *srrm4* splicing and translation-blocking morpholinos were reported to cause a curved body axis and conspicuous deficits in trigeminal neuron neurite arborization (Calarco et al. 2009). We examined the trigeminal ganglion at 30 hpf using an anti-Islet1 antibody to label the cell bodies and acetylated Tubulin to visualize the peripheral axons. In both wild-type (n= 8) and homozygous mutant *srrm4^y712^* (n= 8) embryos, we observed similar arborization, with no salient differences in outgrowth or branching (Figure 7A–B). In addition, we found that *srrm4* mutants from both alleles did not exhibit a curvature of the body axis and were morphologically indistinguishable from wild-type siblings (data not shown). The same morpholinos were used in a subsequent study examining the role of *srrm4* in hair cell development (Nakano et al. 2012). *srrm4* morphants were again reported to have a curved body axis and more than 80% reduction in the number of neuromast hair cells, which could be rescued by injection of wild-type *srrm4* mRNA. Use of G0 crispants has been shown to be an efficient method to screen for phenotypes(Shah et al. 2016; Shankaran et al. 2017; Wu et al. 2018; Kroll et al. 2021); therefore, we generated G0 crispant larvae using gRNAs that targeted both the translation start site and the eMIC domain (targets 4 and 5; Supplementary Figure 2C). Like stable mutants, G0 crispant larvae did not display the previously reported body axis curvature of the morphants (Figure 7C–D). To visualize the hair cells, we stained crispants with FM1-43 and counted hair cell numbers in two lateral line neuromasts per larva. We saw only a small (6%) decrease in hair cell numbers (repeated measures ANOVA, F[1,72] = 9.1, p=0.043 ; Figure 7E), unlike the dramatic reduction reported in morpholino studies. It is unclear why there is a discrepancy between the morphant and mutant phenotypes. It is possible that the morphant phenotypes reflect off-target effects on other genes or that the morphant phenotypes are less apparent in the mutants due to genetic compensation that does not occur with morpholino treatment (reviewed in Arana and Sánchez 2024).

**Figure 7:**
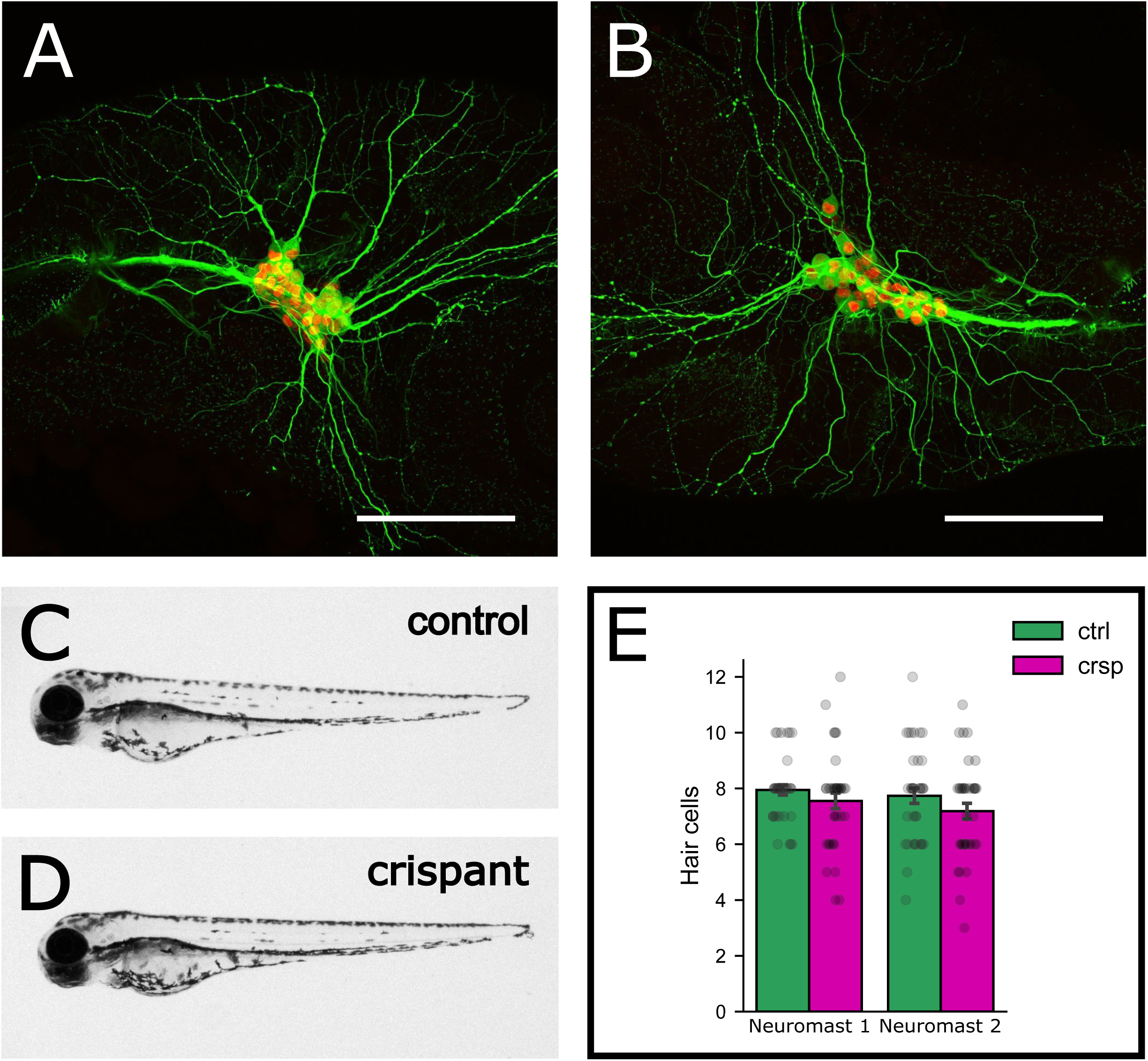
*srrm4* G0 crispants do not recapitulate morphant phenotypes. (A–B) Maximum intensity projections of acetylated Tubulin and Islet 1 antibody staining in trigeminal neurons of wild-type (A) and *srrm4^y712^* homozygous mutants (B) at 30 hpf. Acetylated Tubulin, in green, marks developing neurites and Islet 1, in red, marks trigeminal neuron cell bodies. (C–D) *srrm4* G0 crispant and Cas9-injected control larvae at 3 dpf showing overall body morphology. (C) Cas9-injected larva and (D) *srrm4* G0 crispant larva. (E) Hair cell numbers from two neuromasts per larva in Cas9-injected controls (ctrl) versus *srrm4* G0 crispants (crsp) at 3 dpf. Each dot represents the number of hair cells from an individual neuromast. N = 38 larvae per group.

### *srrm4* crispants have deficits in optic tectum neuropil

To assess whether *srrm4* is required for normal brain development, we utilized whole brain morphometric analyses to look for localized changes in neuronal composition and brain volume in *srrm4* mutants. Our initial analysis utilized G0 crispants injected with multiplexed gRNAs to efficiently screen for phenotypes. Wild-type larvae carrying transgenes that expressed fluorescent proteins throughout the brain were injected with gRNAs that cut in *srrm4* exons 1, 6, and 10. They were then imaged with a confocal microscope at 6 dpf (table 1: crispant experiment 1). We registered the image stacks to reference brains and used CobraZ software to identify regions of the brain with localized changes in neuronal cell types and/or volume (Gupta et al. 2018). Of the thirteen major brain divisions, we found that the optic tectum was significantly smaller in *srrm4* crispants than controls (mean difference = 0.55%, p = 0.016 corrected; Figure 8A). To determine more specifically what regions of the optic tectum were affected, we performed a cluster analysis to identify clusters of contiguous voxels with significant differences in volume (Gupta et al. 2018). We found that all five subdivisions of the optic tectum neuropil were smaller at an uncorrected p < 0.05, and three of the five tectal neuropil regions were still significantly smaller after Holm-Bonferroni correction for multiple region comparisons, further validating the size difference (Figure 8B). To validate the crispant phenotype, we performed 4 additional G0 crispant experiments using non-overlapping gRNAs and different backgrounds for brain registration (Table 1). We then performed a meta-analysis to calculate the effect size for each experiment and a weighted mean based on all five experiments (Figure 8C). Consistent with the original knockdown, the overall effect size indicated a decrease in the size of the optic tectal neuropil in G0 crispants. Interestingly, the only experiment in which no significant decrease was seen was crispant experiment 4, in which a single gRNA was used to target the eMIC domain. This suggests that the tectal neuropil phenotype may be dependent on more N-terminal regions of the Srrm4 protein.

**Figure 8:**
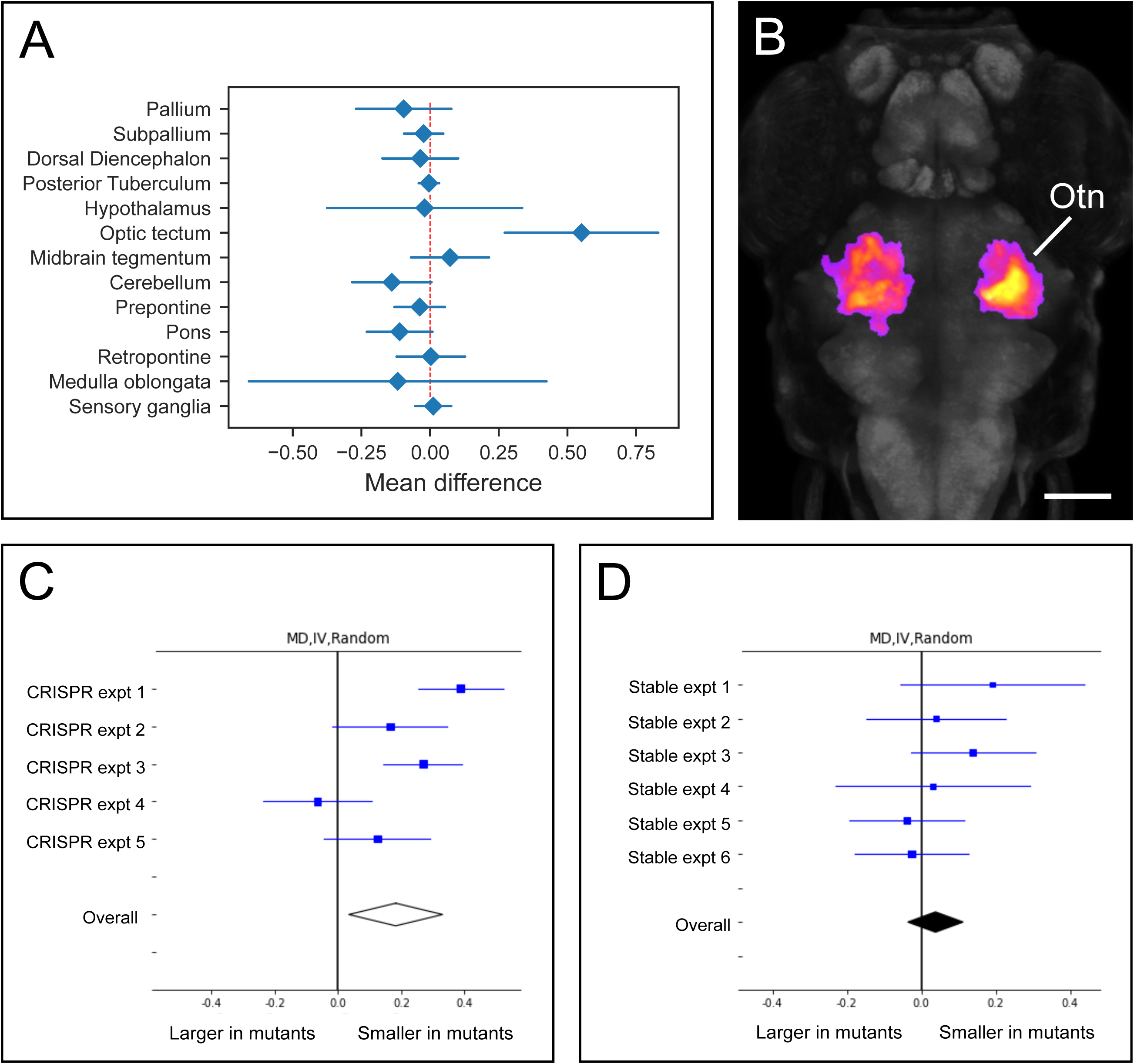
*srrm4* G0 crispants display reduced optic tectum neuropil phenotype not observed in stable mutants. (A) Plot showing mean percent size difference between crispant and control larvae within major brain divisions at 6 dpf. (B) Dorsal view of maximum intensity projection of voxels showing a significant reduction in volume in *srrm4* G0 crispant larvae at 6 dpf based on cluster analysis (fire). Gray staining shows pan-neuronal expression of *elavl3*. Anterior is to the top. Otn: optic tectum neuropil. Scale bar = 100 µm. (C–D) Forest plots depicting random model meta-analyses of *srrm4* morphometry experiments. (C) *srrm4* G0 crispant experiments and (D) and *srrm4* stable mutant experiments. Details of conditions for each experiment provided in Table 1.

Next, we performed the same whole brain morphometric analyses on stable mutants under various conditions (Table 1). Across six experiments, no significant tectal neuropil phenotype was detected, nor was there an effect when we performed a meta-analysis of all experiments, which included “stressing” embryos with Cas9 injections similar to the G0 crispant experiments (experiment 3) and eliminating any potential maternal contribution of *srrm4* (experiment 6; Figure 8D). These data suggest that either the crispant phenotype was due to off target effects, despite reproducibility using multiple non-overlapping gRNAs, or that there was some form of compensation in the stable mutants.

### *srrm4* mutants do not have widespread changes in gene expression levels or alternative exon use

We tested whether alternative splicing of microexons was affected in a *srrm4* predicted loss-of-function mutant. Irimia et al. (2014) reported that in cortical glutamatergic neurons differentiated from mouse embryonic stem cells, the largest changes in microexon splicing occurred at late transitions during differentiation in maturing post-mitotic neurons. Therefore, we sequenced mRNA from *srrm4^y712^* larvae at 3 dpf, when a significant number of neurons are undergoing differentiation and synaptogenesis and analyzed changes in alternative exon usage. We found that when comparing *srrm4^y712^* homozygous mutants to their wild-type siblings, only 4 exons smaller than 27 nt had significant MV[dPSI] scores over 10%, indicating that only a very small number of microexons annotated in the VastDB database were affected by the *srrm4* mutation (Supplementary Table 1).

Given the very subtle effects of *srrm4* mutations on morphological and molecular phenotypes, we asked whether the mutations may be inducing genetic compensation. To do so, we analyzed our RNA-seq data to look for a decrease in *srrm4* expression that would indicate potential nonsense-mediated decay, as well as differences in gene expression that may compensate for reduced *srrm4* function. We found no significant decrease in the expression of *srrm4* and that there were only 7 genes expressed at higher levels in the mutant compared to wild-type at FDR-corrected<0.05 (Supplementary Table 2). None of these genes are predicted to be involved in post-transcriptional processing. We also looked specifically at the expression levels of known splicing factors and did not detect any significant differences in expression between mutant and wild-type samples (Supplementary Table 3). Based on these data, we conclude that genetic compensation through changes in gene expression are unlikely to occur in *srrm4^y712^* mutants.

## Discussion

Microexons are a class of 3–27 nucleotide exons that are highly enriched in neurons and that display distinct temporal expression patterns. However, relatively little is known about the regulation and function of most microexons during neural development. To further our understanding of microexon regulation, we performed a mRNA expression study of *srrm4*, a key regulator of microexon splicing, during early zebrafish brain development and then generated mutant lines for phenotypic analysis.

Our expression analysis revealed that *srrm4* is dynamically expressed in multiple cell types and states in the developing brain. *notch1a* has been used a marker of proliferation zones in the developing zebrafish brain (Mueller and Wullimann 2016). We found that at 2 dpf, *srrm4* co-localizes with *notch1a* in the rhombomeres. Along with *notch1b* and *notch3*, *notch1a* is expressed in rhombomeric centers where it is required for maintenance of a progenitor state (Belmonte-Mateos et al. 2023). From 2 dpf to 5 dpf, *srrm4* showed partial co-localization with *elavl3*, a marker of newly differentiated cells, but was also expressed at high levels in cells not expressing *elavl3*. Consistent with our *in situ* hybridization results, single cell expression data from DanioCell shows a positive correlation of *srrm4* expression with both *notch1a* and *elavl3* in neural tissues (Sur et al. 2023). Thus, our data show that *srrm4* expression is not restricted to a particular developmental state. In the developing hindbrain, we found that *srrm4* partially co- localized with both glutamatergic and glycinergic markers, indicating that *srrm4* expression is also not associated with a particular cell type. Based on these data, *srrm4* function does not appear to be restricted to particular developmental processes, but rather, is likely required for many processes in multiple cell types during neural development.

We focused particular attention on *srrm4* expression in the cerebellum, where *srrm4* and its paralog *srrm3* are expressed at high levels in both mice and zebrafish (Nakano et al. 2019; Shirakawa et al. 2024). In mice, *Srrm3* is strongly expressed in all three layers of the cerebellum, whereas *Srrm4* expression is weak in the Purkinje cells but strong in the molecular and granule layers (Nakano et al. 2019). Consistently, another study reported that mouse *Srrm4* expression in the cerebellum was found to be absent in the GAD67- expressing molecular and Purkinje layers but to overlap with NeuN-expressing granule cells (Shirakawa et al. 2024). Interestingly, zebrafish *srrm3* colocalizes with *srrm4* in granule cell regions but is also expressed in regions of the corpus cerebelli with very little expression of *srrm4*, which we hypothesize may differentiate into Purkinje neurons (data not shown). Based on co-localization with the granule cell marker *vglut1*, we found that *srrm4* is expressed at high levels in the granule cells of both the cerebellum and the torus longitudinalis, a “cerebellar-like” structure that lies dorsal to and along the medial margins of the optic tectum. In addition to *vglut1*, *srrm4* colocalizes with *neuroD1-GFP* and *atoh1c* in the torus longitudinalis (data not shown), suggesting that *srrm4* may be part of a conserved granule cell differentiation program. Given the expression of *srrm3* in these cells, as well, a *srrm3; srrm4* double mutant would likely be needed to uncover the potential role for microexon splicing in granule cell development.

To determine the functions of *srrm4* during brain development, we used CRISPR-based mutagenesis to knock down levels of *srrm4* in G0 animals, as well as to create stable mutant lines for analysis. We first tested previously reported phenotypes for *srrm4* based on morpholino analyses. Unlike in the previous reports, we found that in our genetic knockdowns and mutants, we did not see an abnormal body curvature in the larvae or any other gross morphological defects(Calarco et al. 2009; Nakano et al. 2012), and indeed, homozygous mutants grew to fertile and viable adults. This lack of morphological defects is consistent with that reported by (Ciampi et al. 2022) and (Lopez-Blanch et al. 2024) for a frame-shifting mutation in the eMIC domain of zebrafish *srrm4*. We then tested whether or not our mutants exhibited the trigeminal neurite growth and branching defects reported for *srrm4* morphants (Calarco et al. 2009). At 30 hpf, we did not observe obvious differences in trigeminal neurons between mutants and controls and were, therefore, unable to reproduce the morphant phenotype. (Nakano et al. 2012) used the same morpholino as (Calarco et al. 2009) and reported a reduction in the numbers of neuromast hair cells at 72 hpf, a phenotype that they were able to rescue with injection of wild-type *srrm4* mRNA. We also did not observe the same dramatic reduction in hair cell numbers reported by Nakano et al. (2012) in *srrm4* crispants. As use of CRISPR mutagenesis has increased in recent years, there have been growing numbers of reports of inconsistencies between morphant and mutant phenotypes. Discrepancies may be due to off-target effects of the morpholinos or because of genetic compensation in stable mutant lines (reviewed in Arana and Sánchez 2024). It is unclear why *srrm4* mutants do not recapitulate the morphant phenotypes, but our RNA-seq data did not indicate the likelihood of nonsense-mediated decay-induced genetic compensation through upregulation of other splicing factors. Additionally, we performed our analysis of hair cell numbers using G0 crispants, which are typically not subject to genetic compensation (reviewed in Arana and Sánchez 2024). We therefore hypothesize that the phenotypes previously reported in *srrm4* morpholino knockdowns may, despite careful and rigorous mRNA rescue experiments, be due to off target effects.

As we were unable to reproduce any of the phenotypes previously reported for *srrm4* morphants, we utilized whole brain morphometric analyses to search for mutant phenotypes. This method enables unbiased phenotype discovery, without a priori hypotheses on affected brain regions (Gupta et al. 2018; Bhandiwad et al. 2024). We tested G0 crispants, which are mosaic for mutations in *srrm4*, and stable mutant lines under a variety of conditions. This analysis disclosed a reduction in the size of the optic tectum neuropil in G0 crispants, but not in stable mutants. We first considered whether this was due to off target effects from one or more gRNAs. However, we were able to reproduce the optic tectum phenotype using two non-overlapping sets of gRNAs, so this possibility seems unlikely. It has previously been reported that stable mutant lines do not always display G0 crispant phenotypes due to genetic compensation, in which one or more genes may be upregulated to compensate for the mutation(Buglo et al. 2020). In many cases, genetic compensation appears to be triggered by nonsense-mediated decay (El-Brolosy et al. 2019; Ma et al. 2019). To investigate if genetic compensation may explain the lack of a *srrm4* stable mutant phenotype, we analyzed RNA-seq data from *srrm4^y712^*, the mutant line with an early stop codon that was used for morphometric experiments. We looked for significant changes in gene expression in other zebrafish Ser/Arg repeat genes, notably *srrm3*, which is a paralog of and has overlapping functions with *srrm4*, as well as *srrm1* and *srrm2*. We also looked for changes in other splice factors, as well as in transcriptomic expression overall. We were unable to identify any changes in gene expression that might explain why stable mutants failed to show the same tectal phenotype as crispants. We therefore hypothesize that some other, as yet, unidentified mechanism of compensation may be present.

Previous studies to identify *Srrm4* regulated microexons in mouse focused on a few particular regions of the brain and ear (Quesnel-Vallières et al. 2015; Nakano et al. 2019). To generate a broader and more complete dataset of *srrm4*-regulated microexons, we performed RNA-seq on *srrm4* stable mutant and sibling wild-type larvae at 3 dpf. Surprisingly, we were only able to detect significant changes in splicing of 4 microexons. In addition to the possibility of compensation, we hypothesize that redundancy with *srrm3* may enable largely normal splicing in *srrm4* mutants. However, it is also possible that changes in microexon splicing may be difficult to detect in whole brain samples due to the heterogeneity of cell types and developmental states, as microexon splicing is likely differentially regulated in various neural tissues. Therefore, in order to generate a more complete dataset of *srrm4*-regulated microexon splicing, it may be necessary to perform region-specific RNA-seq at various timepoints during development.

Together, these data show that *srrm4* has a dynamic expression pattern in multiple cell types during neural development, including expression in proliferating and newly differentiated cells. Previous *srrm4* loss-of-function phenotypes using morpholino knockdowns are not reproducible in either *srrm4* crispants or stable mutants, a circumstance that has also been reported in other studies. Rather, *srrm4* mutants do not show an overt morphological developmental deficit, possibly because of an as yet unknown compensation mechanism and/or redundancy with *srrm3*.

## Acknowledgments

This work was supported by the Intramural Research Program of the *Eunice Kennedy Shriver* National Institute for Child Health and Human Development and utilized the high-performance computational capabilities of the Biowulf Linux cluster at the National Institutes of Health, Bethesda, MD. The authors thank Hariom Sharma for designing *glyt2* and *gad1b* oligo probe pools.

