## Supplementary figures and images for "Mutations in the microexon splicing regulator *srrm4* have minor phenotypic effects on zebrafish neural development"

### Supplementary Figures

Supplementary Figure 1

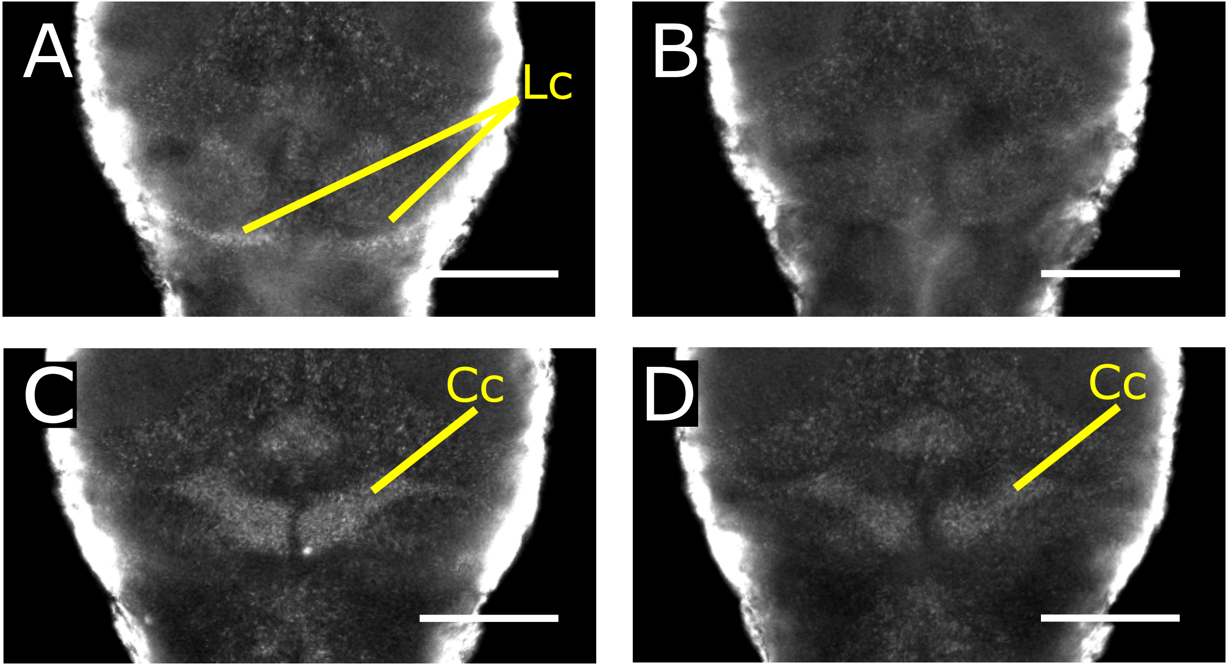

#### Supplementary Figure 2

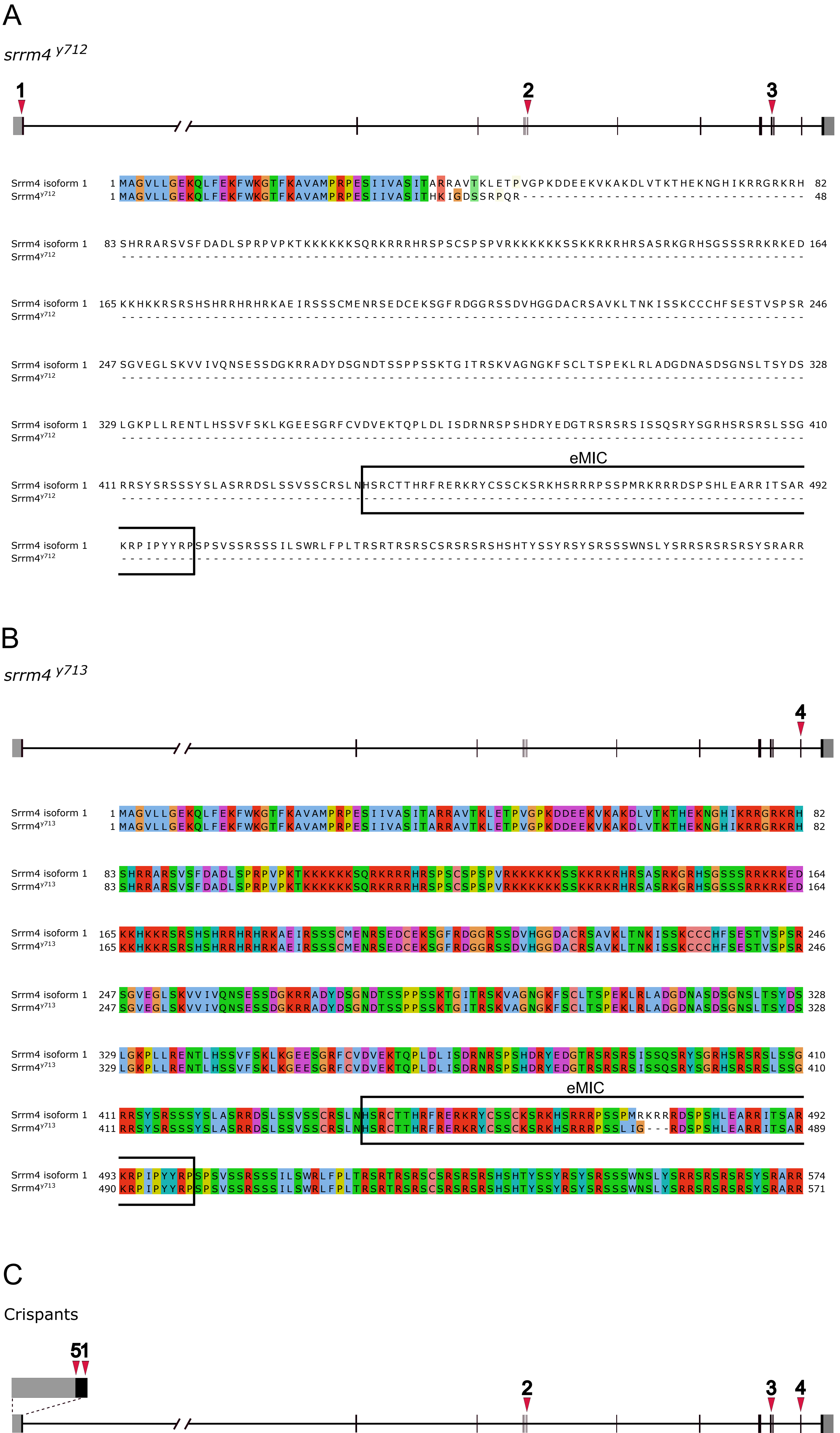
